# Stepwise autophosphorylation regulates biased agonism of the insulin receptor

**DOI:** 10.1101/859868

**Authors:** Na-Oh Yunn, Mangeun Park, Jungeun Noh, Sung Ho Ryu

**Affiliations:** The Department of Life Sciences, Pohang University of Science and Technology, Pohang, 37673, South Korea; School of Interdisciplinary Bioscience and Bioengineering, Pohang University of Science and Technology, Pohang, 37673, South Korea

## Abstract

Insulin is a key regulator of energy metabolism in peripheral tissues but also functions as a growth factor. Insulin binding to the insulin receptor (IR) leads to autophosphorylation of intracellular tyrosine residues, which simultaneously initiates a multitude of signals and functions. In contrast, some artificial (non-insulin) ligands for the IR result in biased agonism, selectively activating the PI3K/AKT pathway and metabolic effects without activating mitogen-activated protein kinase (MAPK) pathway and mitogenic effects. However, the precise mechanism of biased agonism at the receptor level remains unclear. The biased agonist IR-A48 aptamer selectively induces mono-phosphorylation of Tyr1150 residue (m-pY1150) in the kinase domain of IR. Hence, we hypothesized that IR autophosphorylation is a stepwise process in which formation of m-pY1150 represents an intermediate step. To explore this idea, we used hybrid receptors in which insulin can bind only one of the two binding sites of the dimeric receptor. Asymmetric insulin binding selectively induced symmetric m-pY1150 in both kinase domains. Moreover, the juxtamembrane domain, which interacts with the kinase domain, restricted the full activation of IR, and symmetric m-pY1150 played a crucial role in the rearrangement of intracellular domains to release this restriction. Our findings demonstrate that the symmetry of insulin binding to a dimeric receptor determines the stepwise autophosphorylation of IR. Furthermore, considering that the degree of ligand symmetry on a receptor depends mainly on ligand concentration, our results suggest that IR may play metabolic-biased roles in peripheral tissues dependent on local insulin concentrations.

## Introduction & Results

The insulin receptor (IR), a member of the receptor tyrosine kinase (RTK) family, exists as a homo-dimeric receptor containing two extracellular α-subunits and two transmembrane β-subunits linked by disulfide bonds (α2β2)^1^. Insulin secreted from pancreatic beta cells binds to the ectodomain of IR on the plasma membrane, leading to autophosphorylation of intracellular tyrosine residues^2^. Activation of IR simultaneously initiates two major signaling pathways, the phosphoinositide 3-kinase(PI3K)/AKT pathway and the mitogen-activated protein kinase (MAPK) pathways. The metabolic effects of insulin, such as glucose uptake and lipid synthesis, are mainly regulated by the PI3K/AKT pathway^3^. By contrast, the MAPK pathway is the primary mediator of mitogenic effects of insulin, although the PI3K/AKT pathway is also involved^4,5^.

Surface receptors are commonly depicted as on/off switches that propagate intracellular signaling following ligand binding to the receptor. However, the discovery of biased agonism (also known as functional selectivity), in which a ligand selectively initiates certain signaling pathways, has revealed that receptor activation is more complicated than simple switches, and that receptors can be primary determinants of functional regulation by ligands^6^. Some non-insulin IR ligands based on peptides, antibodies, and aptamers can selectively activate the PI3K/AKT pathway and its metabolic effects, but in contrast to insulin, these agents have weaker mitogenic effects^7^. Over the past decades, biochemical and structural studies on IR have suggested several plausible models for insulin-mediated IR activation^8-10^. However, these canonical models for IR activation cannot explain how the biased agonists regulate receptor autophosphorylation and intracellular signaling in a distinct manner from insulin. In particular, IR-A48, an aptamer agonist of IR, not only selectively activates the PI3K/AKT pathway and glucose uptake, but also induces mono-phosphorylation of Tyr1150 residue (m-pY1150) in the kinase domain of IR without phosphorylating other tyrosine residues^11^. This led us to hypothesize that m-pY1150 is a key link between the biased intracellular signaling and receptor modulation by biased agonists.

### m-pY1150 is a key characteristic of biased agonists to the IR

To investigate the relationship between m-pY1150 and biased agonism, we examined site-specific phosphorylations of IR by another biased agonist. S519 peptide is an insulin mimetic consisting of an N-terminal 20-residue peptide (S519-N20) and a C-terminal 16-residue peptide (S519-C16); a derivative of S519 exerts biased agonism to induce AKT phosphorylation and metabolic effects similar to IR-A48^12-14^. To assess whether S519 also induces m-pY1150, we administered S519-N20, S519-C16, and S519 to Rat-1 cells overexpressing human IR (Rat-1/hIR) and analyzed the site-specific tyrosine phosphorylation of IR. As expected, S519-C16 selectively induced m-pY1150 similarly to IR-A48, whereas S519-N20 had no effect on IR autophosphorylation (Fig. 1a). The role of S519-N20 is to increase the efficacy of S519-C16: S519 induced significantly higher levels of m-pY1150 than S519-C16, even at a 10-fold lower concentration.

**Figure 1:**
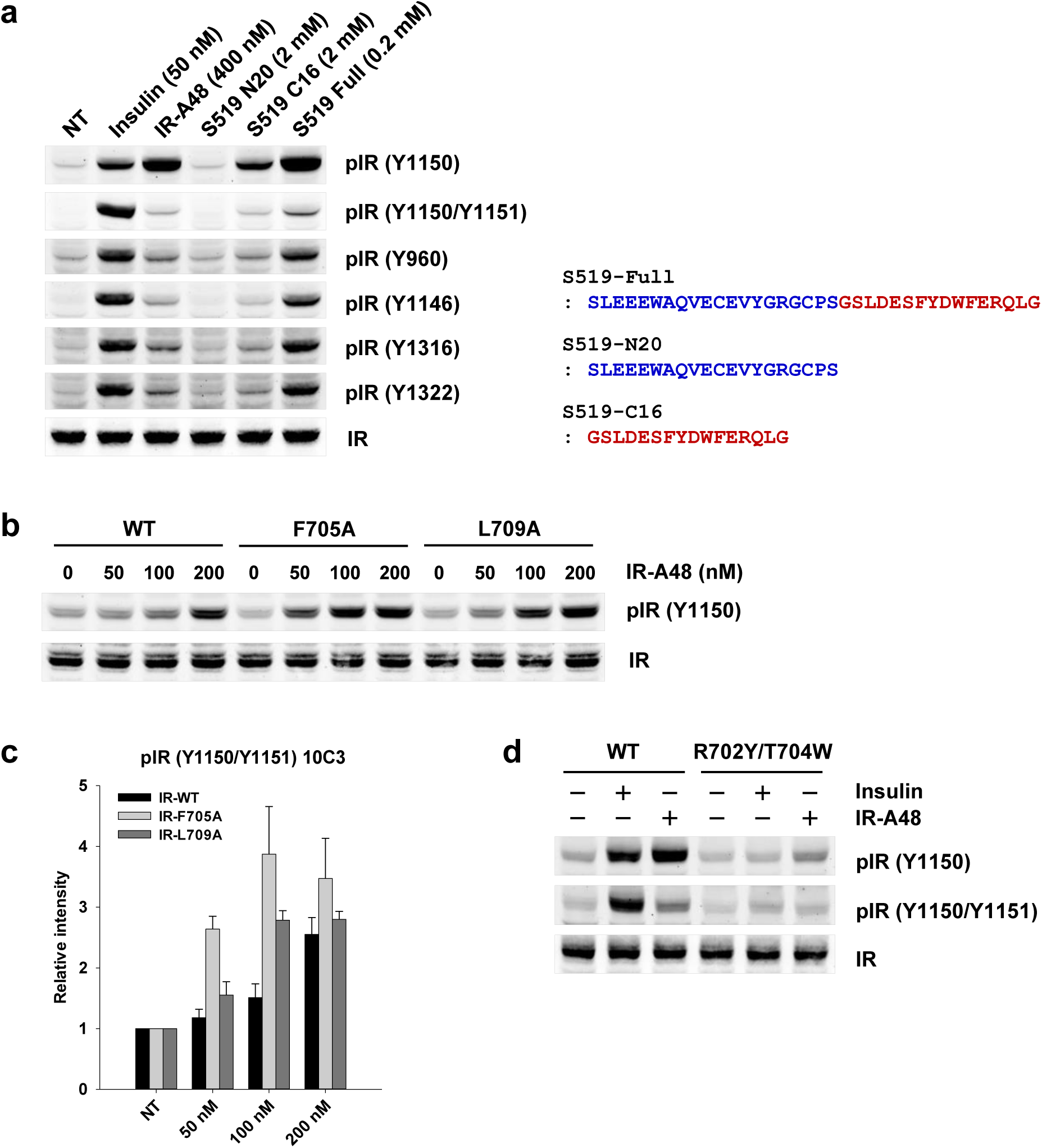
m-pY1150 is a key characteristic of biased agonists of insulin receptor. **a**, Serum-starved Rat-1/hIR cells were stimulated with insulin for 5 min, IR-A48 aptamer for 1 hour, or peptides for 1 hour. **b**, CHO-K1 cells were transiently transfected with wild-type IR (WT), IR with the Phe705Ala mutation (F705A), or IR with the Leu709Ala mutation (L709A). After serum starvation, the cells were stimulated with IR-A48 aptamer for 1 hour. **c**, Quantification of phosphorylation levels in **(b)**. Graph shows mean ± S.E. of three independent experiments; data are normalized against the band intensity of the corresponding negative control (NT) and thus express fold change relative to basal. **d**, CHO-K1 cells were transiently transfected with wild-type IR (WT) or IR with the Arg702Tyr (R702Y) and Thr704Trp mutations (T704W). Cells were stimulated with 200 nM insulin for 10 min or 1 μM IR-A48 aptamer for 1 hour.

A structural analysis of S519-C16 revealed that it binds at exactly the same position as the helical C-terminal (αCT) segment on the surface of the first leucine-rich repeat (L1) domain, demonstrating that the insulin-mimetic activity of S519 arises from competitive displacement of αCT^15^. Thus, we introduced an Ala substitution at the Phe705 (F705A) or Leu709 (L709A) residue of αCT to disrupt the interaction between αCT and L1, and then introduced both Tyr and Trp substitutions in the Arg702 and Thr704 residues, respectively (R702Y/T704W), to strengthen the interaction^16,17^. The F705A and L709A mutations obviously increased induction of m-pY1150 by IR-A48 relative to wild-type (WT) IR, whereas the R702Y/T704W mutation completely inhibited IR-A48 activity (Fig. 1b,c,d). These results indicate that m-pY1150 is a key characteristic of biased agonists for IR, and that displacement of the αCT segment from L1 domain is a critical step in induction of m-pY1150.

### Interaction between Fn3-1 and CT segment is critical for IR autophosphorylation

The fact that two different ligands were selective for m-pY1150 implies that formation of m-pY1150 is a natural process of IR autophosphorylation caused by an intrinsic mechanism, which is specifically activated by biased agonists. Hence, we hypothesized that m-pY1150 represents an intermediate step between basal state and full activation state in insulin-mediated IR activation. Previous studies suggested that insulin binds first to the primary binding site (site 1) composed of the L1-αCT complex, and then forms a crosslink to the secondary binding site (site 2) located in the first fibronectin type III-1 domain (Fn3-1)^10,18,19^. Moreover, the αCT segment undergoes significant displacement on the surface of L1 when insulin engages with site 1 of IR^16^. Therefore, our first hypothetical model was that insulin binding to site 1 specifically induces m-pY1150, and that the crosslink to site 2 induces full IR autophosphorylation (Extended Data Fig. 1a). To test this model, we performed alanine scanning of eight residues (P495, D496, F497, R498, R539, S540, N541, E575) in the Fn3-1 domain, which is expected to contain site 2 based on a cryo-EM analysis of the IR-insulin complex (PDB:6CE9)^19^. Alanine substitution of only two residues (D496A and R498A) significantly disrupted both m-pY1150 and pY1150/pY1151, but no m-pY1150 selectivity was observed (Fig. 2a). To rule out the possibility that insulin binding to other residues complemented the effect of a single Ala substitution, we introduced dual-Ala substitutions of the five residues (P495, R539, S540, N541 and E575) except for hydrophobic residue F497, but decrease in phosphorylation of IR was not observed (Extended Data Fig. 1b). These results indicate that that these five residues do not engage in functional interactions with insulin.

**Figure 2:**
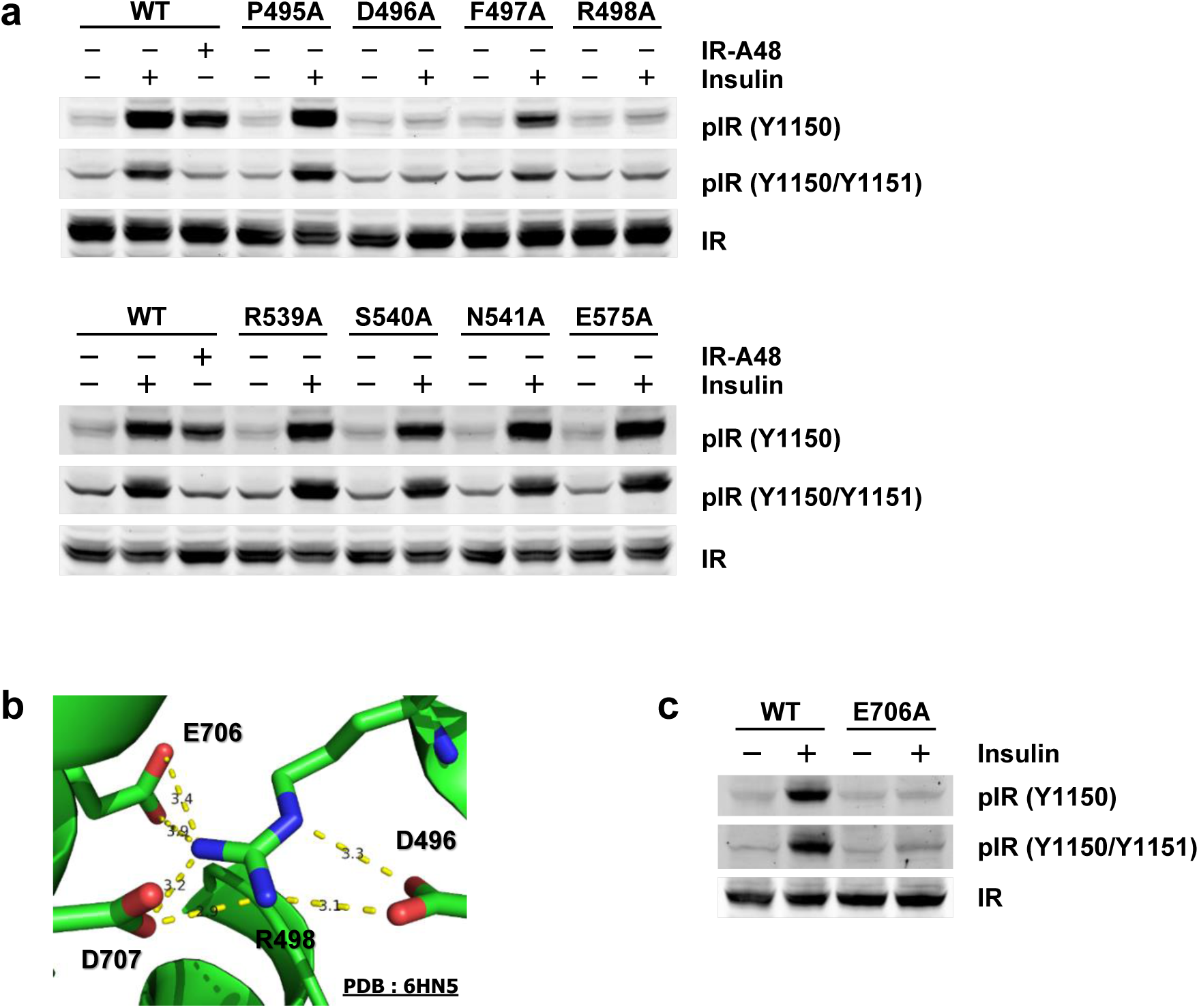
Interaction between Fn3-1 and CT segment is critical for IR autophosphorylation. **a**, CHO-K1 cells were transiently transfected with wild-type IR (WT) or the indicated IR mutant. After serum starvation for 1 hour, the cells were stimulated with 200 nM insulin for 10 min or 1 μM IR-A48 aptamer for 1 hour. **b**, Atomic interactions among Asp496, Arg498, Glu706, and Asp707 in the complex of insulin and the ectodomain of IR (PDB:6HN5)<u>^18^</u>. **c**, CHO-K1 cells were transiently transfected with wild-type IR (WT) or IR harboring the Glu704Ala mutation. After serum starvation for 1 hour, the cells were stimulated with 200 nM insulin for 10 min.

Interestingly, another cryo-EM analysis of the IR-insulin complex (PDB:6HN5) revealed that D496 and R498 potentially form electrostatic interactions with Glu706 and Asp707 in the αCT segment, but not with insulin (Fig. 2b)^18^. Consistent with the structural analysis, alanine substitution of Glu706 (E706A) completely disrupted insulin-mediated IR autophosphorylation, even though Glu706 is not involved in insulin binding to site 1 of IR (Fig. 2c)^16^. These results conflicted with our first hypothetical model, but demonstrated that the interaction between the Fn3-1 and CT segment, rather than insulin binding to Fn3-1, is important for insulin-mediated IR autophosphorylation (Extended Data Fig. 1c). These observations indicate that the conception of site 2 as the secondary site of insulin binding to IR must be revised.

### Asymmetric insulin binding selectively induces m-pY1150

Our second model to explain the intermediate step involved the ligand symmetry of a dimeric receptor. Because IR exists in a dimeric form linked by disulfide bonds, holo-IR has two symmetric insulin-binding sites. However, insulin binding exhibits negative cooperativity, and only one insulin asymmetrically binds to a holo-receptor at physiological concentrations^10^. Hence, we hypothesized that the first asymmetric insulin binding to IR induces m-pY1150, and that binding of the second insulin to the other site fully activates IR autophosphorylation (Fig. 3a). To test this theory, we introduced an Ala substitution of residue Phe64 (F64A) into one side of a dimeric receptor to perturb insulin binding asymmetrically^16,20^. In addition, we conjugated YFP to the C-terminus of one of the monomers to distinguish the phosphorylation state of each monomer of the dimeric receptor. Insulin induced phosphorylation of YFP-conjugated IR normally, whereas symmetric F64A clearly disturbed this process. However, in the WT+F64A/YFP hybrid receptor, asymmetric insulin binding induced m-pY1150 in YFP-conjugated IR to the same degree as in WT, but phosphorylation of other Tyr residues was significantly reduced (Fig. 3b). This result suggests that asymmetric binding of the first insulin to a dimeric receptor selectively induces m-pY1150, and that binding of the second insulin to the other site initiates full phosphorylation of other tyrosine residues.

**Figure 3:**
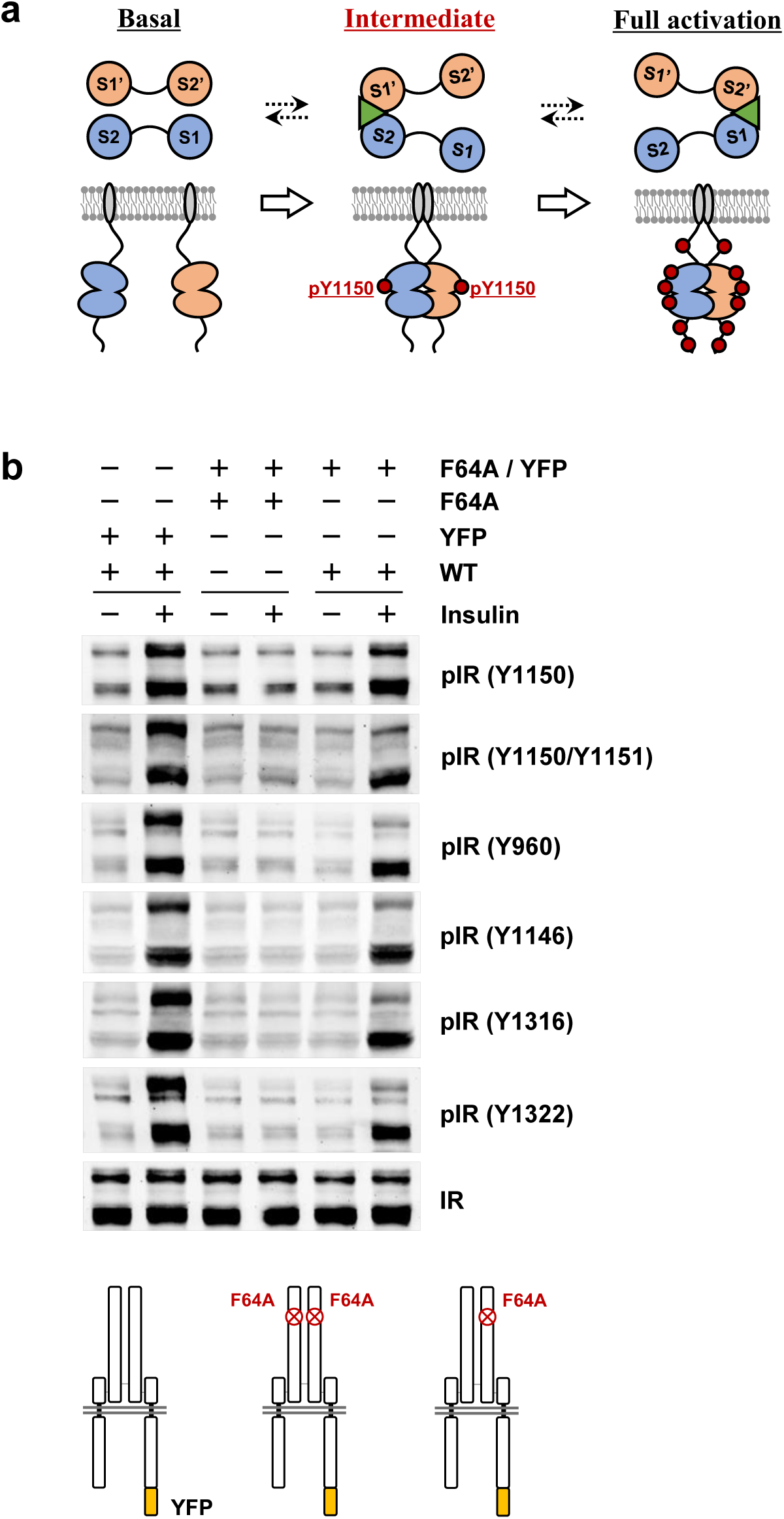
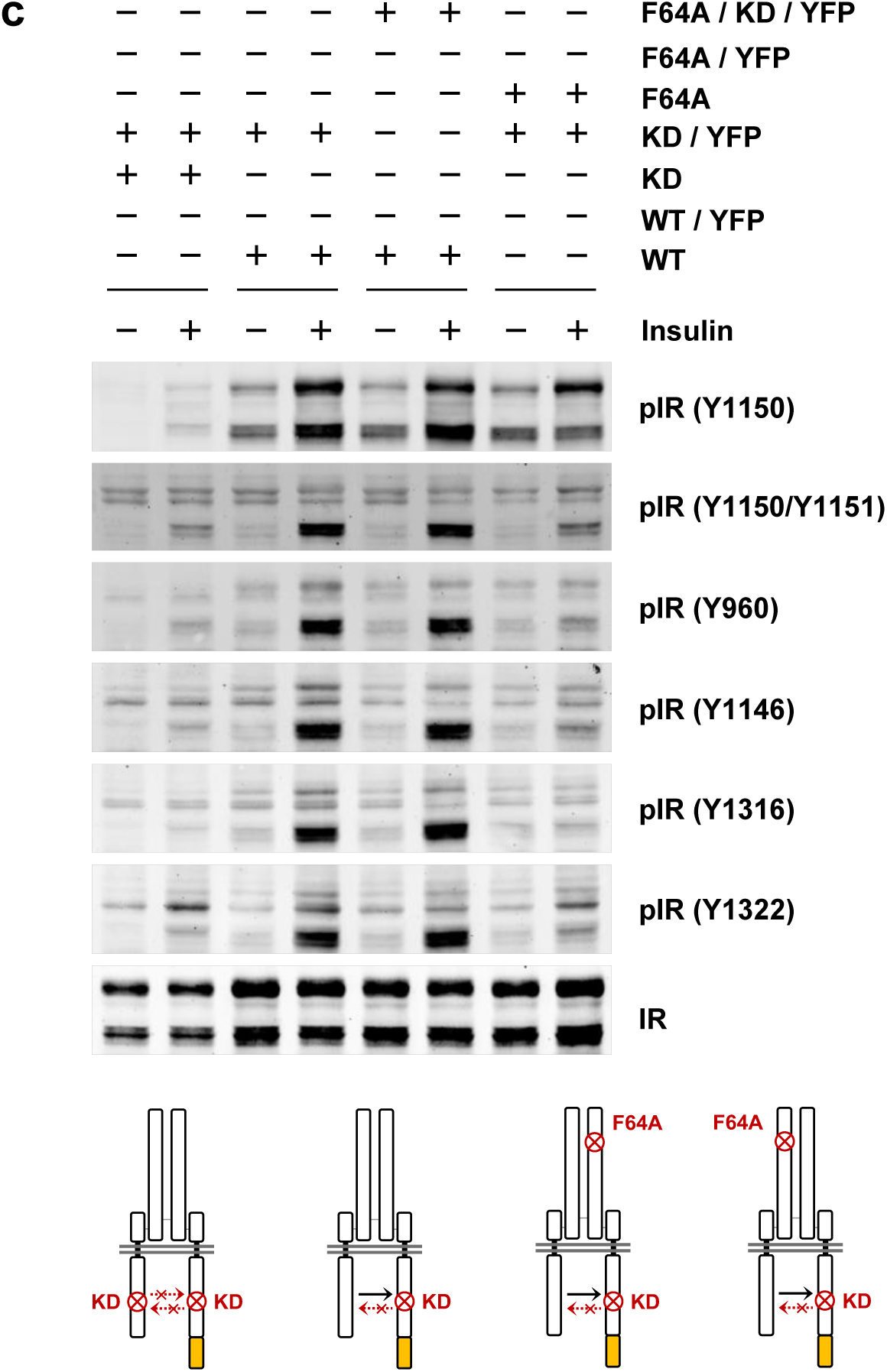
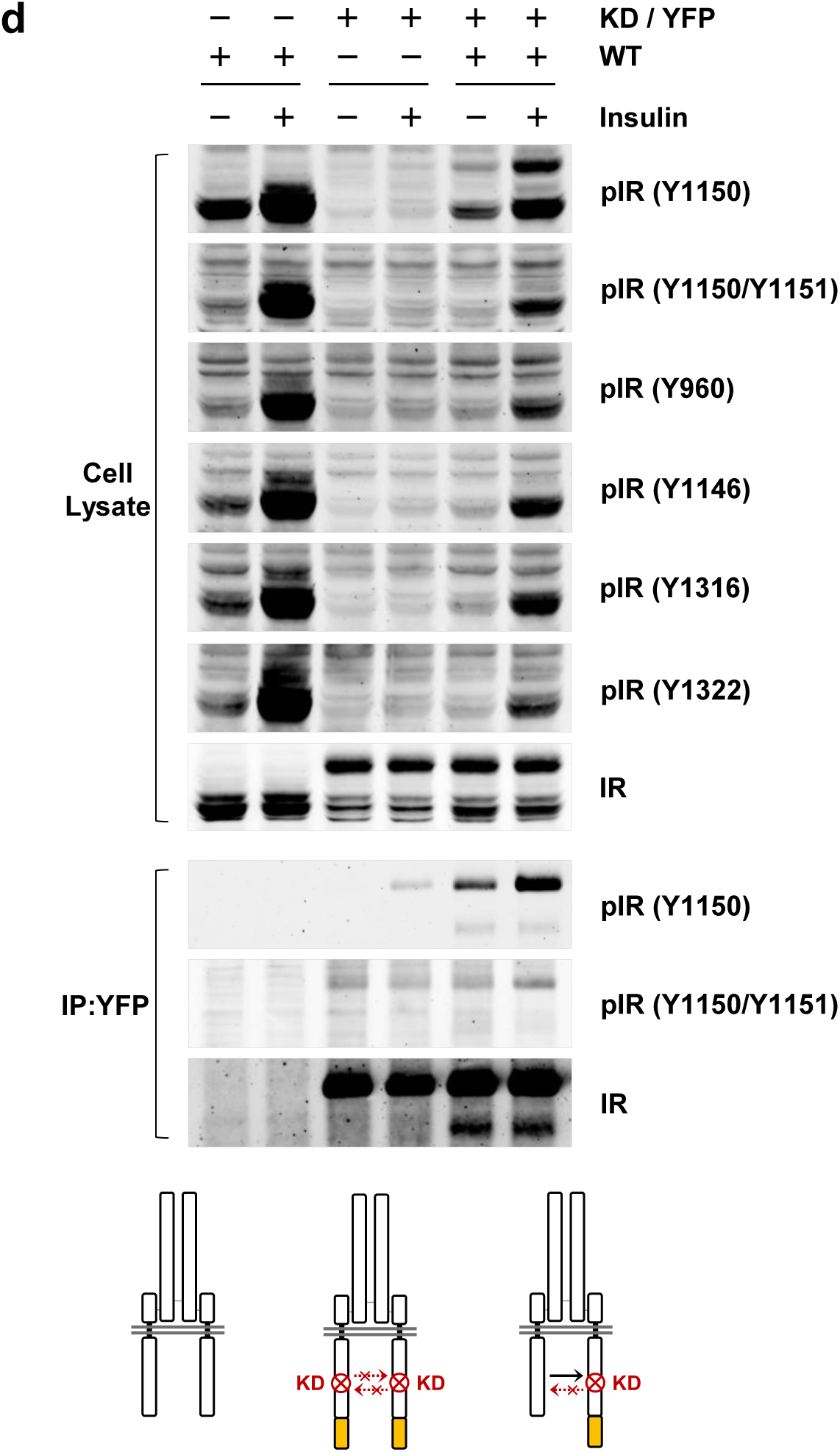
Ligand symmetry and symmetric m-pY1150 determine full autophosphorylation of IR. **a**, Hypothetical model of ligand symmetry. **b, c, d**, To generate hybrid IRs, CHO-K1 cells were transiently co-transfected with short-IR (without YFP, lower bands) and long-IR (with YFP, upper bands; DNA ratio = 1:2). After serum starvation for 1 hour, the cells were stimulated with 200 nM insulin for 10 min. **WT**=wild type, **F64A**=Phe64Ala mutation, **YFP**=yellow fluorescent protein, **KD**=kinase-dead mutation.

Because IR autophosphorylation occurs in *trans*, we wondered whether asymmetric insulin binding determines the direction of *trans*-autophosphorylation of m-pY1150. To address this question, we introduced a kinase-dead (KD, K1018A) mutation into YFP-conjugated IR to allow *trans*-autophosphorylation to occur in only one direction, and changed the location of the asymmetric F64A. Regardless of the direction of asymmetric binding, insulin induced m-pY1150 of IR harboring the KD mutation at a similar level, indicating that asymmetric insulin binding induces m-pY1150 symmetrically in both kinases of a dimeric receptor (Fig. 3c). Unexpectedly, only m-pY1150 was observed in the WT+KD/YFP hybrid receptor, even though both insulin-binding sites were normal. We reproduced this experiment and performed immunoprecipitation of YFP to rule out the possibility of non-specific detection. As before, WT kinase coupled with a KD partner kinase induced only m-pY1150 in *trans*, demonstrating that symmetric m-pY1150 of a dimeric receptor is significantly involved in the full activation of IR (Fig. 3d).

### The juxtamembrane domain plays a key role in the restriction of full activation

These results suggest two characteristics of IR autophosphorylation. First, symmetric m-pY1150 in a dimeric receptor is an essential step for the full activation of IR. Second, m-pY1150 determines the substrate specificity of IR kinase (Extended Data Fig. 2a). Substrate specificity of kinases is dependent on the amino-acid sequences of substrates (peptide specificity), or regulated by the frequency of encounter between kinase and substrate (substrate recruitment)^21^. To verify that the change in peptide specificity of the IR kinase was dependent on phosphorylation status, we performed an *in vitro* kinase assay using partially purified IR. Basal IR following serum starvation and activated IR following insulin stimulation were immunoprecipitated from Rat-1/hIR, and each purified IR was incubated with a substrate peptide containing the IR activation loop sequence in the presence of ATP. The level of m-pY1150 was higher in activated IR kinase than in basal IR kinase, but basal IR kinase also induced m-pY1150 normally. However, pY1150/pY1151 was not detected in either IR kinase (Fig. 4a). Because dual phosphorylation of a substrate requires that it contact the kinase twice, we also performed an *in vitro* kinase assay using a substrate containing m-pY1150. As before, pY1150/pY1151 was still not detected, indicating that phosphorylation of the activation loop increases the activity of IR kinase, but does not change peptide specificity.

**Figure 4:**
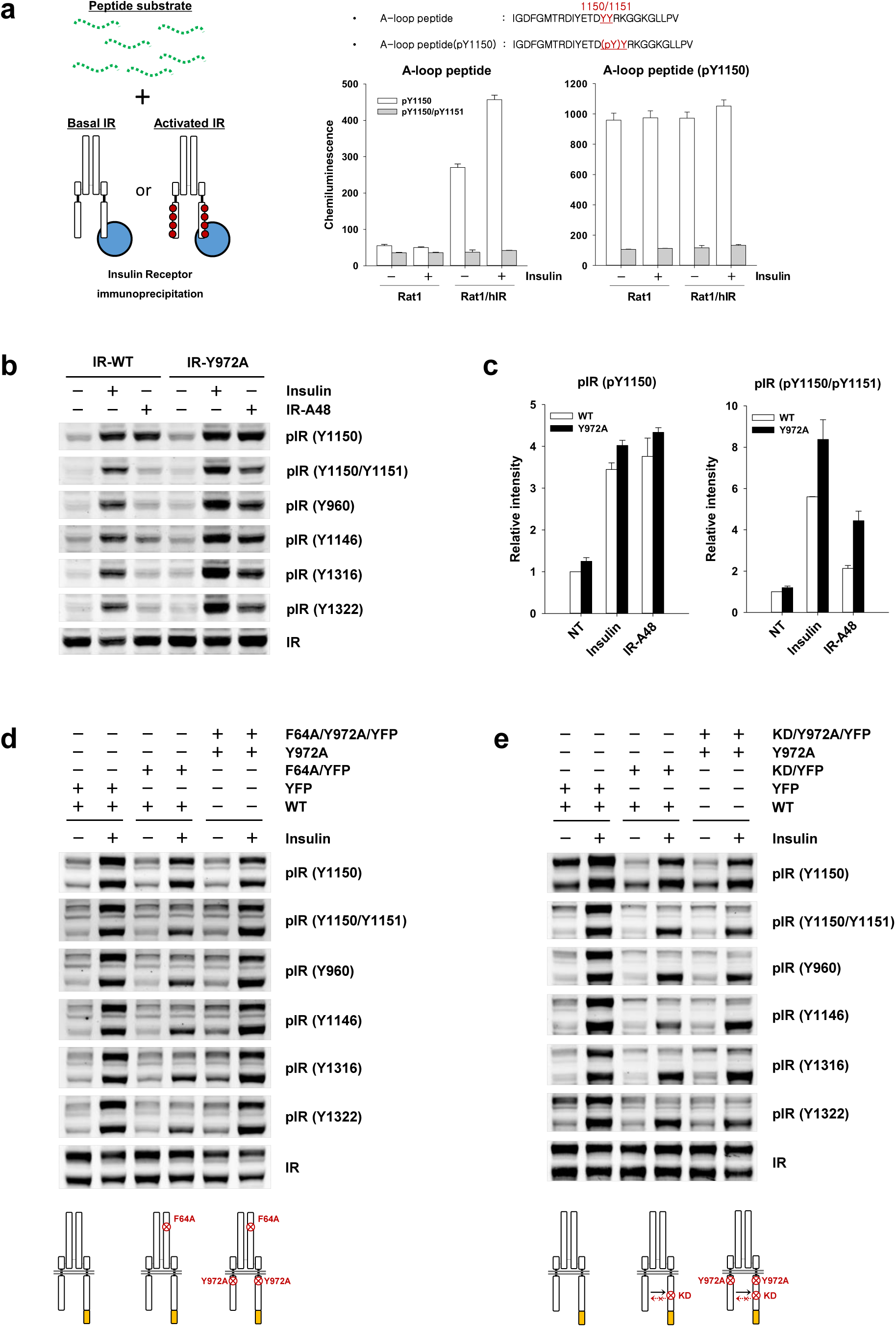
The juxtamembrane domain plays a key role in restriction of full activation. **a**, *In vitro* kinase assay for IR kinase. IR was partially purified from Rat-1/hIR cells by immunoprecipitation. Purified Irs were incubated with biotin-conjugated A-loop peptides for 60 min at 36°C. After peptide immobilization on streptavidin-coated 96-well plates, phosphorylation was detected using site-specific primary antibody and alkaline phosphatase-conjugated secondary antibody. **b, d, e**, To generate hybrid IR, CHO-K1 cells were transiently co-transfected with each short IR (without YFP) and long-IR (with YFP; DNA ratio = 1:2). After serum starvation for 1 hour, the cells were stimulated with 200 nM insulin for 10 min. **WT**=wild type, **F64A**=Phe64Ala mutation, **Y972A**=Tyr972Ala mutation, **YFP**=yellow fluorescent protein, **KD**=kinase-dead mutation. **c**, Quantification of phosphorylation levels of **(b)**. Graph shows means ± S.E. of three independent experiments, and data are normalized against the band intensity of the corresponding negative control (NT), and thus show fold change relative to basal.

Therefore, we speculated that the role of m-pY1150 in the substrate specificity of IR kinase is related to substrate recruitment. In many RTKs, the juxtamembrane domain plays a *cis*-autoinhibitory role in receptor autophosphorylation. In particular, residue Tyr972 on the juxtamembrane domain of IR represses its kinase activity in the basal state by interacting with the N-terminal lobe of the kinase domain^22^. Alanine substitution of Tyr972 (Y972A) not only increases catalytic efficiency (∼4-fold) relative to WT, but also leads to a significant increase in basal phosphorylation of IR (∼30-fold). Similarly, alanine substitution of Asp1149 (D1149A), which plays a critical role in blocking substrate access to the active site of IR kinase, increases catalytic efficiency (∼10-fold) even more strongly than Y972A. However, in contrast to catalytic efficiency, D1149A has a modest effect (∼3-fold) on basal phosphorylation of IR relative to Y972A^22,23^. The difference between the two mutations in basal phosphorylation of the receptor and catalytic efficiency suggests that Tyr972 also plays an important role in conformational rearrangement of β-subunits, allowing *trans*-autophosphorylation of the receptor. Thus, we verified the m-pY1150 selectivity of IR-A48 aptamer in the Y972A mutant. In contrast to WT, IR-A48 aptamer induced phosphorylation of all Tyr residues, similarly to insulin, in the Y972A mutant (Fig. 4b,c). Moreover, the Y972A mutation also restored phosphorylation of Tyr residues by insulin in the WT+F64A/YFP hybrid receptor (Fig. 4d). However, the WT+KD/YFP hybrid receptor always formed m-pY1150 in response to insulin regardless of the Y972A mutation, indicating that symmetric m-pY1150 in a dimeric receptor is required for the release of the juxtamembrane domain (Fig. 4e). Taken together, our observations show that the interaction between the juxtamembrane domain and the kinase domain prevents Tyr residues other than Tyr1150 from being a substrate for *trans*-autophosphorylation, and that the release of the juxtamembrane domain induced by symmetric m-pY1150 leads to a conformational rearrangement of β-subunits, allowing IR kinase to phosphorylate other Tyr residues (Extended Data Fig. 2b).

## Discussion

Our findings demonstrate that autophosphorylation of IR is a stepwise process regulated by ligand symmetry of the dimeric receptor (Extended Data Fig. 2c). The binding of the first (asymmetric) insulin to IR results in only symmetric m-pY1150 in both kinases. In this intermediate step, the insulin receptor exhibits biased agonism, selectively activating the PI3K/AKT pathway and metabolic effects such as glucose uptake. Moreover, symmetric m-pY1150 plays an essential role in the release of the juxtamembrane domain from the kinase domain, allowing phosphorylation of other Tyr residues. Binding of the second insulin to the opposite site of a dimeric receptor fully activates the insulin receptor (IR), inducing the MAPK pathway and mitogenic effects. One insulin molecule binds IR with picomolar affinity (0.2–0.5 nM)^24-26^. Because the level of insulin in the blood ranges from 0.04 nM (after overnight fasting) to 0.2 nM (after meal) under normal physiological condition, we anticipate that most IRs in peripheral tissues remain in the intermediate state, which is biased toward metabolic effects.

From physiological insulin concentrations to 100 nM, only one insulin molecule binds a dimeric receptor with high affinity^10,18^. The binding of two or more insulins to a dimeric receptor occurs at much high insulin concentration, but phosphorylation of all Tyr residues is clearly observed at physiological insulin concentrations <10 nM. Moreover, cryo-EM analysis of the insulin and IR complex revealed that insulin binds independently to canonical site 2 located between the Fn3-1 and Fn3-2 domains; thus a maximum of four insulins can form a complex with a dimeric receptor^27^. However, we confirmed that m-pY1150 selectivity is not dependent on insulin binding to canonical site 2 (Extended Data Fig. 3a). Therefore, the binding of two or more insulins to a receptor does not seem to be necessary for full activation of the receptor.

An unanswered question in this model is how binding of the first and second insulin binding results in different phosphorylation status in a dimeric receptor. Considering only the ectodomain, the active conformations of the receptor induced by the first and second insulin are identical^18^. One plausible explanation for this is that the binding of each insulin results in asymmetric rearrangement of intracellular domains. For example, activation of epidermal growth factor receptor (EGFR) is mediated by the formation of an asymmetric dimer of kinase domains^28^. However, because the kinase domains of IR and insulin-like growth factor 1 receptor form symmetric dimers during *trans*-autophosphorylation and in full activation state, we expected that rearrangement of the intracellular domains of IR is different from EGFR^29,30^. It remains to be elucidated how the conformational changes in the ectodomain induced by insulin binding are transmitted to rearrangement of intracellular domain to initiate downstream signaling. Accordingly, structural analysis of full-length IR in the basal, intermediate, and active states is necessary for a deeper understanding.

Biased agonism has been intensively studied in G protein-coupled receptors^6^. However, very few cases have been reported in RTKs, and the underlying mechanism remains unclear^7^. Most RTKs share conserved mechanisms for ligand-induced receptor activation, including receptor dimerization, *trans*-autophosphorylation, and *cis*-autoinhibition by the juxtamembrane domain and activation loop in the kinase domain^31^. In light of our findings, it is reasonable to assume that the mechanism of biased agonism of IR could be generalized to other RTKs. Thus, our work may shed light on the functional modulation of RTKs dependent on local ligand concentration and receptor density on the plasma membrane.

## Acknowledgment

This research was supported by the Global Research Laboratory (GRL) Program through the National Research Foundation of Korea (NRF) funded by the Ministry of Science and ICT (No. NRF-2016K1A1A2912722).

## Materials and Methods

### Reagents and antibodies

Aptamer was synthesized by Aptamer Science (Pohang, Korea). Bovine insulin (I5500) was purchased from Sigma-Aldrich (St. Louis, MO, USA). Phospho-peptides conjugated to biotin for ELISA assay were synthesized by GenScript (Hong Kong). Antibodies against the IR β-subunit (sc-57342) and phospho-IR (pY1150, sc-81500), and IR β-subunit antibody-conjugated agarose beads (sc-57342 AC) were purchased from Santa Cruz Biotechnology (Dallas, TX, USA). Antibodies against phospho-IR (Y960, 44-800G), phospho-IR (Y1150/Y1151, 44-804G), phospho-IR (Y1316, 44-807G), and phospho-IR (Y1322, 44-809G), as well as alkaline phosphatase (AP)-labeled anti-rabbit/mouse antibodies and CSPD substrate for AP, were purchased from Invitrogen (Carlsbad, CA, USA). Phospho-IR (Y1146, 3021) antibody was purchased from Cell Signaling Technology (Danvers, MA, USA). IRdye 680LT– conjugated anti-rabbit/mouse antibodies were purchased from LI-COR (Lincoln, NE, USA).

### Cell culture and experiments

Rat-1 cells overexpressing human IR (Rat-1/hIR) were maintained in high-glucose Dulbecco’s modified Eagle’s medium (DMEM) with 10% (vol/vol) fetal bovine serum (FBS; Gibco). CHO-K1 was maintained in Ham’s F-12K medium with 10% (vol/vol) FBS. All cells were incubated at 37°C under a humidified atmosphere containing 5% CO_2_ prior to experiments. Before insulin or aptamer stimulation, cells were incubated in serum-free media for 1 hr for serum starvation and then incubated in Krebs-Ringer HEPES buffer (25 mM HEPES pH 7.4, 120 mM NaCl, 5 mM KCl, 1.2 mM MgSO4, 1.3 mM CaCl_2_, 1.3 Mm KH2PO4) for 1 hour. For cell experiments, aptamer and insulin were prepared in Krebs-Ringer HEPES buffer. All aptamer samples were heated for 5 min at 95°C and slowly cooled at room temperature to reconstitute the tertiary structure of aptamer. Aptamer and insulin samples were warmed to 37°C before experiments.

### Western blot and immunoprecipitation

To prepare total cell lysate, harvested cells were lysed in buffer containing 50 mM Tris-HCl (pH 7.4), 150 mM NaCl, 1 mM EDTA, 20 mM NaF, 10 mM β-glycerophosphate, 2 mM Na_3_VO_4_, 1 mM PMSF, 10% glycerol, 1% Triton X-100. Soluble cell lysate was isolated by centrifugation at 14,000 rpm for 15 min at 4°C. For immunoprecipitation, isolated cell lysate was incubated with 2 μg antibodies overnight at 4°C, and protein A agarose bead was added to pull down antibodies. After SDS-PAGE, transferred membranes were incubated with primary antibodies overnight at 4°C. Blots were visualized using the LI-COR Odyssey infrared imaging system.

### IR kinase assay

Rat-1/hIR or Rat-1 cells were lysed in buffer containing 50 mM Tris-HCl (pH 7.4), 150 mM NaCl, 1 mM EDTA, 20 mM NaF, 10 mM β-glycerophosphate, 2 mM Na_3_VO_4_, 1 mM PMSF, 10% glycerol, 1% Triton X-100, and protease inhibitor cocktail. Soluble cell lysate was isolated by centrifugation at 14,000 rpm for 15 min at 4°C. One milliliter of cell lysate (1 μg/μL) was incubated with 50 μL of IR β-subunit antibody-conjugated agarose bead overnight at 4°C to partially purify IR. The beads were washed three times with lysis buffer, washed twice times with kinase assay buffer (50 mM HEPES pH 7.4, 150 mM NaCl, 10 mM MgCl_2_, 2 mM MnCl_2_, 2 mM Na_3_VO_4_, 0.05% Triton X-100), and finally resuspended in 800 μL kinase assay buffer. ATP (200 μM) and biotin-conjugated peptide substrates were diluted in kinase assay buffer, and 150 μL bead suspension was mixed with 150 μL substrate/ATP solution. Mixtures were mixed for 60 min at 36°C with vortexing (1000 rpm). After incubation, the samples were boiled for 10 min at 36°C to inactivate IR kinases. A 100 μL aliquot of supernatant was transferred to streptavidin-coated 96-well plates and incubated for 1 hour at room temperature. Synthetic phospho-peptides (20 pmol/100 μL) overnight at 4°C were covalently cross-linked to 96-well plates (Costar) coated with N-oxysuccinimide ester in PBS. After blocking using PBS + 1% BSA for 1 hour RT, the plates were washed three time in TTBS buffer (50 mM Tris-HCl pH 7.6, 150 mM NaCl, 0.05% Tween-20). Primary antibodies diluted 1:1000 in TTBS were incubated for 1 hour at RT and washed three times using TTBS. Next, AP-conjugated secondary antibodies diluted 1:2000 were incubated for 1 hour at RT and washed three times in TTBS. CSPD substrate for AP was added (100 μL per well), and chemiluminescence was measured using a luminometer (Luminoskan Ascent).

---

**Extended Data Figure 1.**
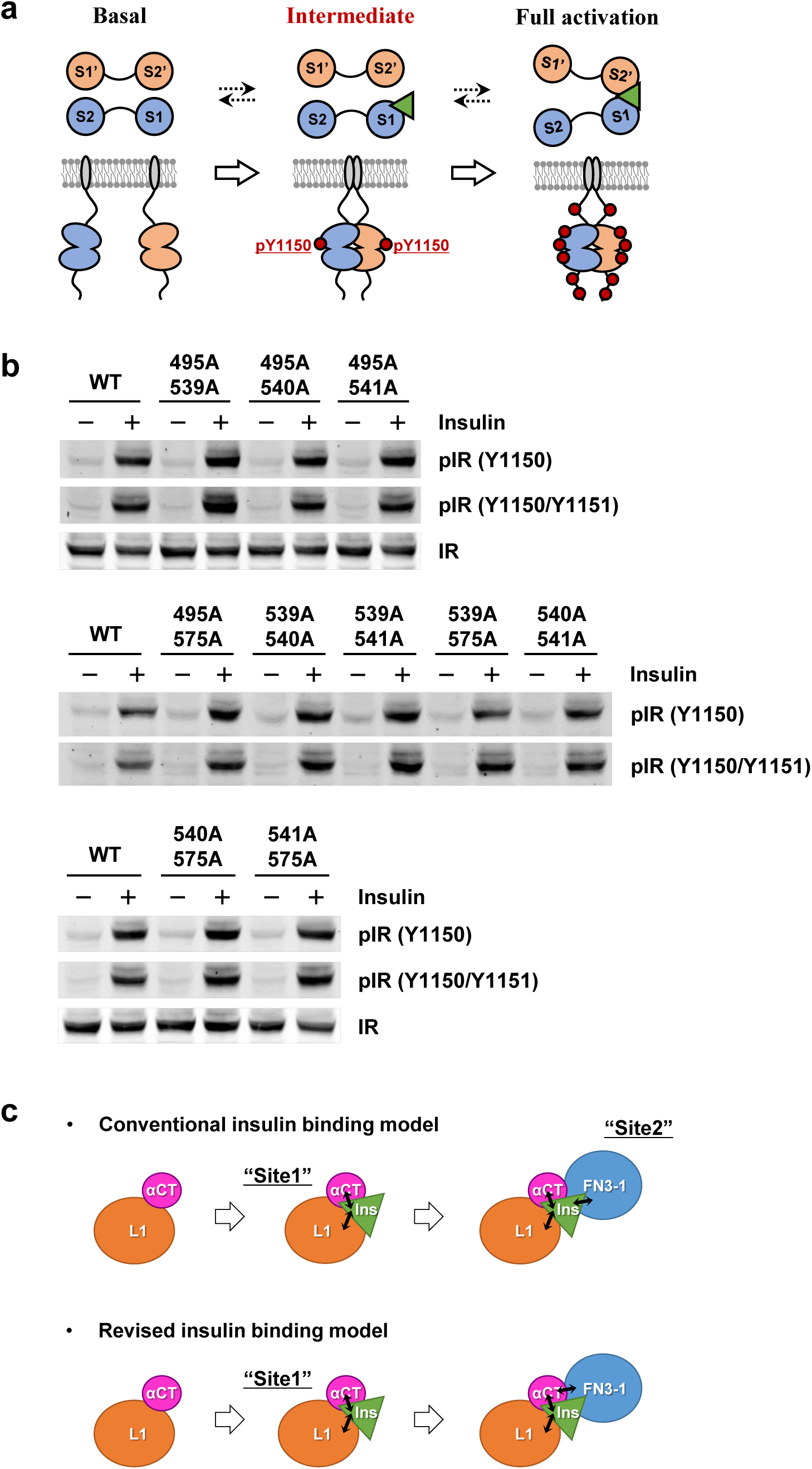

**Extended Data Figure 2.**
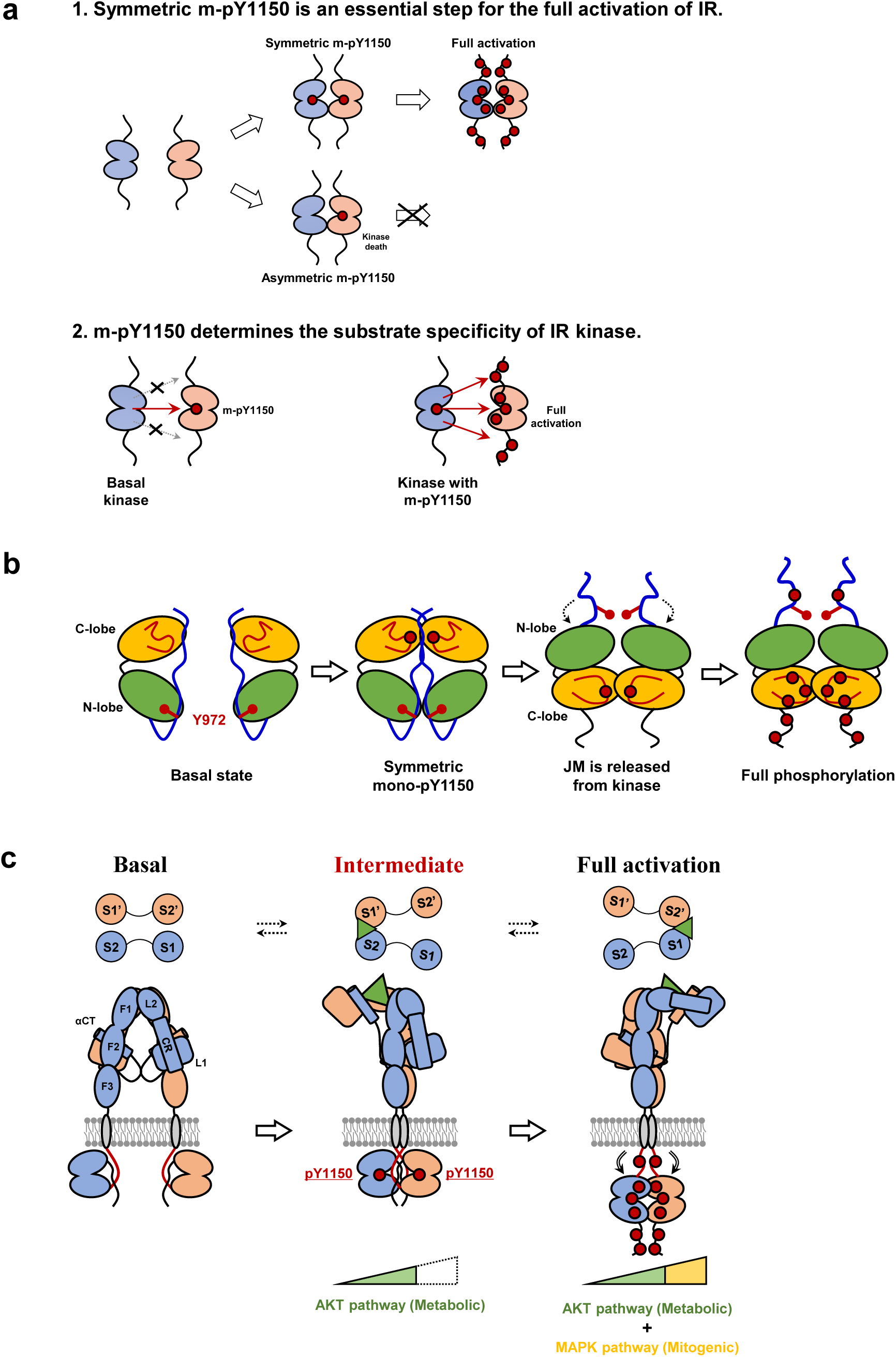

**Extended Data Figure 3.**
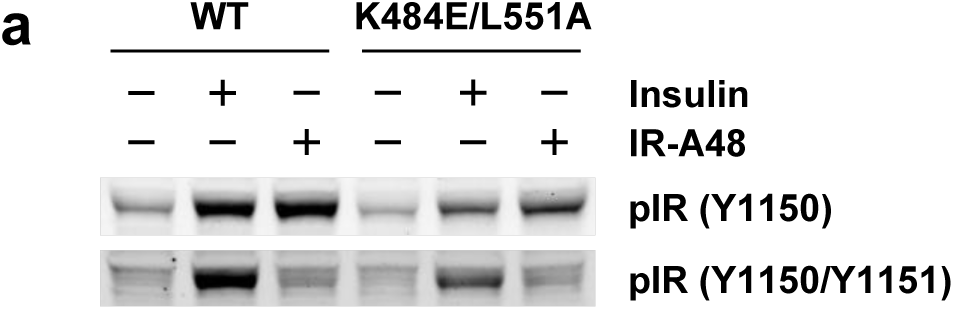

